# The *Drosophila* PDGF/VEGF signaling pathway regulates host immunometabolism in response to parasitoid infection

**DOI:** 10.1101/2025.10.01.679819

**Authors:** Ashley L. Waring-Sparks, Abraham Y. Kpirikai, Jun Yang, Riitvek Baddireddi, Najeeb Marun, Nicholas M. Bretz, Hayden Gosnell, Humayl Malik, David A. Hendrix, Nathan T. Mortimer

## Abstract

Mounting an immune response requires energy, but how that energy is reallocated at the organismal level remains poorly understood. In *Drosophila melanogaster*, infection by a parasitoid wasp triggers a systemic metabolic switch known as immunometabolism which is characterized by a shift in metabolic activity and the redistribution of resources away from organismal development and toward the production of a cellular immune response. We identify the PDGF/VEGF (PVF) signaling pathway as a key initiator of this immunometabolic switch. Genetic manipulation of PVF signaling alters infection outcomes, modulates systemic metabolite profiles, and reveals a direct trade-off between immune function and development. Our findings establish the *Drosophila-*parasitoid wasp system as a genetically tractable model for understanding the molecular basis of immunometabolism.

## INTRODUCTION

Organisms commonly encounter potential pathogens in their environment, and in response to this exposure, mount a defensive immune reaction. Host immune responses have evolved to protect against an array of pathogens, such as macroparasites and microbes, and can include both cellular and humoral mechanisms. Interestingly, alongside these characteristic immune responses, pathogen infection also leads to changes in host metabolism ^1^. These immune-induced changes in metabolism, also known as immunometabolism, have long been recognized and are observed in a wide range of species, including plants, mammals, and invertebrates such as *Drosophila melanogaster* ^1–5^. Immunometabolic changes are highly consistent within species, and progress predictably following pathogen infection, even before clinical signs or symptoms of infection appear ^2,6,7^. Recent findings suggest that immunometabolic changes serve a dual purpose: they can both be a consequence of the reallocation of limited resources to fuel the immune response ^4,8^ or they can act as direct mediators of immunity via roles in immune signaling or pathogen killing and clearance ^9^.

Mounting an immune response is energetically costly for the infected host and necessitates a systemic metabolic adaptation ^10,11^. Classic life-history theory predicts that the immunometabolic switch is indicative of a reallocation of the energy and nutrient resources necessary to mount an immune response ^1^. This theory is based on the premise that organisms have finite resources, including stored cellular energy or metabolite pools, that are simultaneously required for distinct physiological and behavioral functions; thus when an organism invests in one process, it limits the resources available to other processes ^12^. This reallocation leads to trade-offs which can be seen in the antagonistic relationships between the immune response and reproduction success or development rate in a variety of host species ^13–15^. The observed plasticity in resource allocation allows organisms to respond to changing conditions, including pathogen infection ^1^.

Despite the undisputed need for energy reallocation for immune responses, it is becoming increasingly appreciated that metabolites play additional important roles in the response to infection. It has been proposed that the immunometabolic switch following infection may act as an immune defense in its own right, by depriving pathogens of required nutrients or through the production of immunometabolites that function in immune signaling or that act as cytotoxic molecules in pathogen killing mechanisms ^16–18^. The metabolic switch following infection therefore likely plays multiple important roles in the host immune defense. However, the genetic mechanism that initiates the switch to altered metabolism following infection is largely unclear ^19^. While specific alterations in host metabolic pathways can be accomplished by immune-induced alterations in gene expression, changes in hormone signaling, or altered regulation and function of metabolic enzymes ^2,20–22^, the upstream regulators of these processes remain incompletely understood.

*Drosophila melanogaster* is a widely used genetic model organism to understand basic biological processes, including the mechanisms underlying innate immunity ^23^. The *D. melanogaster* immune system relies on both humoral mechanisms, such as the production of antimicrobial peptides (AMPs) in the fat body via the Toll and Imd NF-κB pathways, and cellular mechanisms involving distinct hemocyte (immune cell) responses such as phagocytosis, coagulation, and the encapsulation of macroparasites ^23–26^. Previous work has demonstrated that mounting an immune response in *D. melanogaster* comes at a cost to other physiological processes, such as reproduction or development, highlighting the conservation of classic life-history trade-offs ^8,27,28^. This suggests that *D. melanogaster* immunity may also allow us to uncover the genetic mechanisms driving immunometabolic reallocation.

Parasitoid wasps are a common natural pathogen of *D. melanogaster* that lay their eggs and transfer venom proteins directly into the host hemocoel during infection. These parasitoids comprise a large and diverse group of species, and among the *D. melanogaster*-infecting parasitoid wasps, species are classified as either virulent and capable of suppressing host immune defenses to ensure successful parasitization (*e.g. Ganaspis hookeri and Leptopilina heterotoma*) or avirulent and susceptible to host encapsulation and elimination (*e.g. Leptopilina clavipes*) ^29–31^. This variation in virulence makes infection of *D. melanogaster* by parasitoid wasps a powerful model of an innate immune response against an ecologically relevant pathogen ^32^. Following parasitoid infection, *D. melanogaster* mounts an encapsulation response that is mediated by the sequential action of immune cell (hemocyte) subtypes. This response is initiated by a conserved calcium-dependent activation of plasmatocytes, a macrophage-like hemocyte subtype, and subsequent activity of immune signaling pathways such as Toll and JAK-STAT ^29,31,33–36^. As in mammalian systems, this immune activation is associated with the production of distinct families of cytokines including the Unpaired cytokine family, the Platelet-Derived Growth Factor/Vascular Endothelial Growth Factor-related proteins (PVFs), Spätzle, Eiger, Growth Blocking Peptide (GBP), and Edin ^37^.

In the case of an avirulent wasp infection response, circulating plasmatocytes are activated via calcium signaling and migrate towards and surround the invading wasp egg forming an inner plasmatocyte layer ^30,31^. Infection also stimulates the production of a distinct hemocyte class, the lamellocyte, which adheres to the plasmatocyte layer, forming an impermeable capsule that is subsequently melanized, leading to parasitoid death ^30,38^. This encapsulation response is energetically costly, and due to the reallocation of metabolic resources, larvae that resist parasitoid infection often show reduced fecundity and smaller body size, which lowers their competitive ability for resources ^27,39,40^. However, if the immune response fails or is suppressed by the venom of a virulent wasp species, the wasp egg continues to develop into a larva that will consume and kill the fly host. So, faced with energetic demands of mounting an immune defense or succumbing to parasitism, *D. melanogaster* larvae shift their metabolism, redirecting resources from growth and development toward immunity to improve their chances of survival ^4,41^.

Recent work reveals that carbohydrate metabolism is a central player in these parasitoid infection-mediated trade-offs. In *D. melanogaster*, the disaccharide trehalose serves as the primary energy source ^42^, and is released from glycogen stores in the fat body and muscle ^4,43^. Upon parasitoid infection, hemocytes upregulate the expression of *Trehalase* (*Treh*) and the trehalose transporter *Tret1-1*. Elevated levels of Tret1-1 facilitate the increased uptake of trehalose which is then metabolized by Treh into glucose and subsequently glucose 6-phosphate (G6P), a key intermediate that feeds into both glycolysis and the pentose phosphate pathway (PPP)^44^. The enzyme *glucose-6-phosphate dehydrogenase* (G6PD) is transcriptionally upregulated following infection ^29^ and this increased G6PD activity directs G6P into the oxidative branch of the PPP. This pathway generates critical metabolites such as nicotinamide adenine dinucleotide phosphate (NADPH) and ribose 5-phosphate (R5P) ^45^. NADPH is vital for fatty acid synthesis, which is required for membrane remodeling during lamellocyte differentiation ^44^. Concurrently, the generated R5P supports de novo nucleotide synthesis, which is essential for the rapid proliferation of immune cells and the transcriptional changes during an immune response ^44^. This switch to PPP activation is associated with increased fly survival following parasitoid infection ^44^.

These, and other recent findings suggest that Drosophila hemocytes are a good model for the role of mammalian immune cells in metabolism and metabolic-related disease ^8^. We have used the *Drosophila*-parasitoid wasp system as a model to study the genetic basis of the immunometabolic switch, and uncovered a role for the *Platelet-Derived Growth Factor/Vascular Endothelial Growth Factor-related* (*PDGF/VEGF*) *factor 1* (*Pvf1*) cytokine and *PDGF/VEGF receptor* (*Pvr*) signaling in initiating the immunometabolic switch. The PVF pathway has been previously linked to developmental metabolism in *D. melanogaster* ^46,47^, and we find that the PVF pathway is also active in metabolic reallocation following infection. Genetic manipulation of the PVF signaling pathway in immune cells alters the outcome of parasitoid infection and influences fly development. More specifically, loss of PVF signaling leads to reduced immune function whereas constitutive activation of the pathway in hemocytes leads to global metabolite changes indicative of a precocious metabolic switch in the absence of infection. Flies in which the PVF pathway is constitutively activated fail to complete development, suggesting that PVF signaling is a potential genetic mechanism for the immune response: development trade-off.

## RESULTS

### PVF signaling is required for hemocyte-mediated encapsulation in the *Drosophila* immune response

The *D. melanogaster* encapsulation response to parasitoid infection is dependent on cytokine mediated signal transduction. PVF family cytokines have been previously linked to *D. melanogaster* metabolism in other contexts ^46–48^, and the *Pvf factor 1* (*Pvf1*) is transcriptionally upregulated following parasitoid infection ^29^, leading us to hypothesize that PVF signaling may play a role in the metabolic reallocation following infection. We predicted that if PVF signaling is linked to immunometabolism, then loss of PVF activity would lead to decreased immune competence. To test this prediction, we expressed a dominant-negative form of the PVF receptor *Pvr* (*UAS-Pvr.DN*^D1^)^49^ in specific classes of hemocytes, and assayed the ability of larvae with impaired PVF signaling to encapsulate eggs from the avirulent wasp *Leptopilina clavipes*. We expressed the *UAS-Pvr.DN^D1^* construct in circulating plasmatocytes with *eater-Gal4* (abbreviated *eater>Pvr.DN*) ^50^ and in immune-activated plasmatocytes and lamellocytes with *msn-Gal4* (abbreviated *msn>Pvr.DN*) ^51,52^. Larvae were dissected 72 hours after being infected by *Leptopilina clavipes*, and we found that both *eater>Pvr.DN* (Figure 1A; p = 1.31x10^-7^) and *msn>Pvr.DN* (Figure 1B; p = 5.05x10^-6^) significantly impaired larval encapsulation ability when compared to genetic background controls.

**Figure 1.**
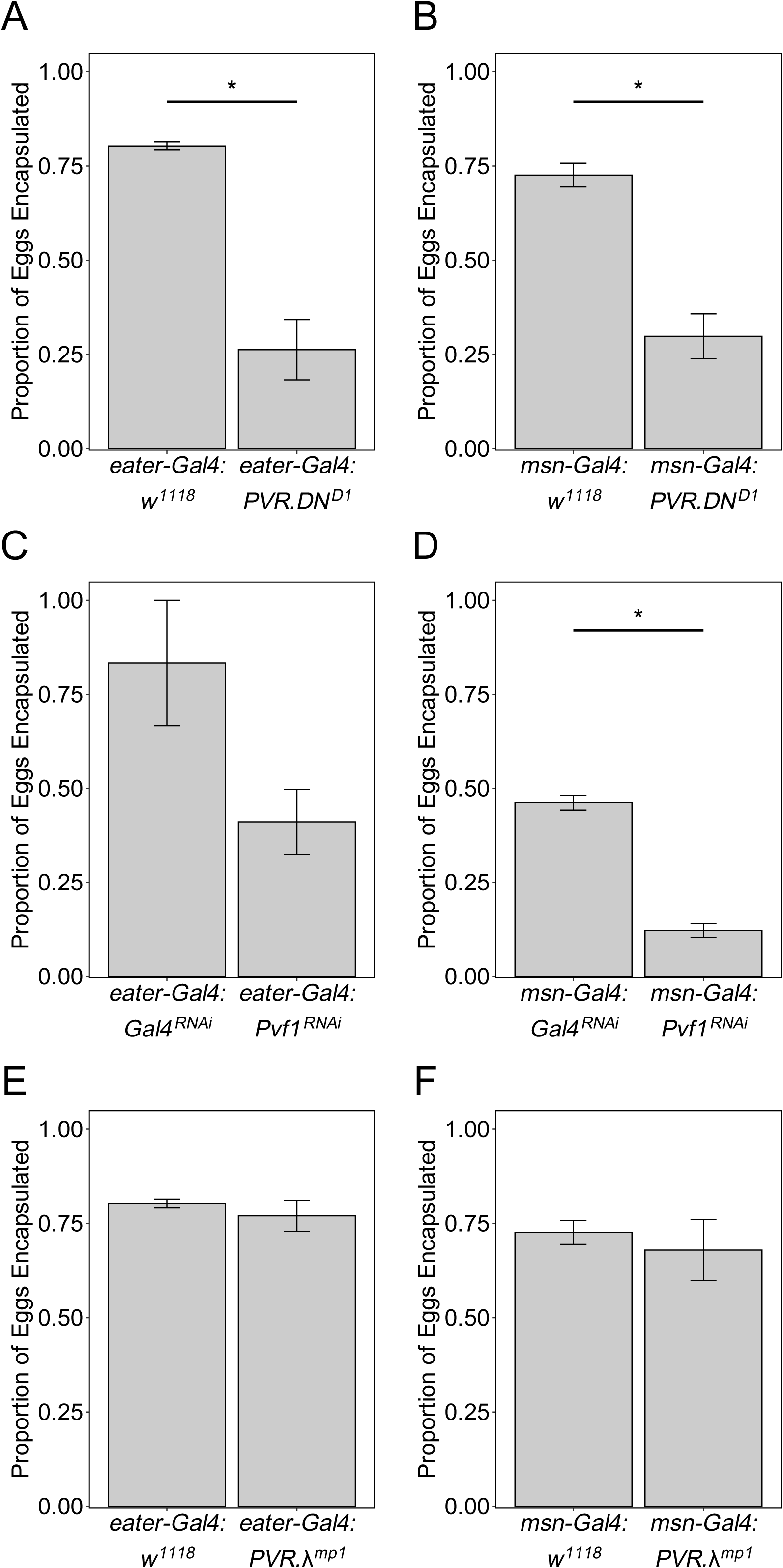
The PVF signaling pathway plays a role in the *Drosophila* cellular immune response. Comparison of the proportion of parasitoid eggs encapsulated in the indicated genotypes. * indicates p < 0.05.

The *Pvf1* cytokine is induced following wasp infection ^29^, and to confirm our results we tested whether tissue specific RNA-interference (RNAi) mediated knockdown of *Pvf1* (*UAS-Pvf1^RNAi^*) could impair the encapsulation response in comparison with a *UAS-GAL4^RNAi^* control. Interestingly, we found that *eater>Pvf1^RNAi^* did not impact encapsulation success (Figure 1C; p = 0.145), whereas *msn>Pvf1^RNAi^* significantly decreased encapsulation ability compared to control (Figure 1D; p = 3.52x10^-4^). These findings suggest that PVF signaling is required in both plasmatocyte and lamellocyte immune cell subtypes for the encapsulation response, and that the signal is initiated by *Pvf1* production specifically in immune activated plasmatocytes or lamellocytes.

To test whether ectopic activation of PVF signaling impacts the cellular immune response, we expressed a constitutively active form of *Pvr* (*UAS-Pvr.λ^mp1^*) ^49^. We found that neither *eater>Pvr.λ* (Figure 1E; p = 0.809) nor *msn>Pvr.λ* (Figure 1F; p = 0.484) had an impact on encapsulation success in comparison with genetic background controls. This suggests that PVF activity is not limiting for the encapsulation of *L. clavipes*.

### Global metabolomics identifies distinct temporal immunometabolic signatures following infection

To test whether PVF signaling plays a role in metabolic reallocation, we first need to better define the immunometabolic state. Previous studies have focused on a narrow range of metabolites, but to better understand *D. melanogaster* immunometabolism, we characterized global metabolic changes between naïve and *L. clavipes* infected larvae at two time points following infection (Figure 2A). The first (or ‘early’) time point is 6-9 hours post infection (hpi), which corresponds to the immune activation phase of the encapsulation response ^30,31^. The second (or ‘late’) time point is 18-21 hpi, which corresponds to a point in the encapsulation response during which the capsule has begun forming on the wasp egg, and there is maximal lamellocyte production ^30,38,52^. We performed metabolomic analysis using avirulent wasp infection in order to isolate immune-induced metabolic changes from potential metabolic effects of parasitoid venom proteins.

**Figure 2.**
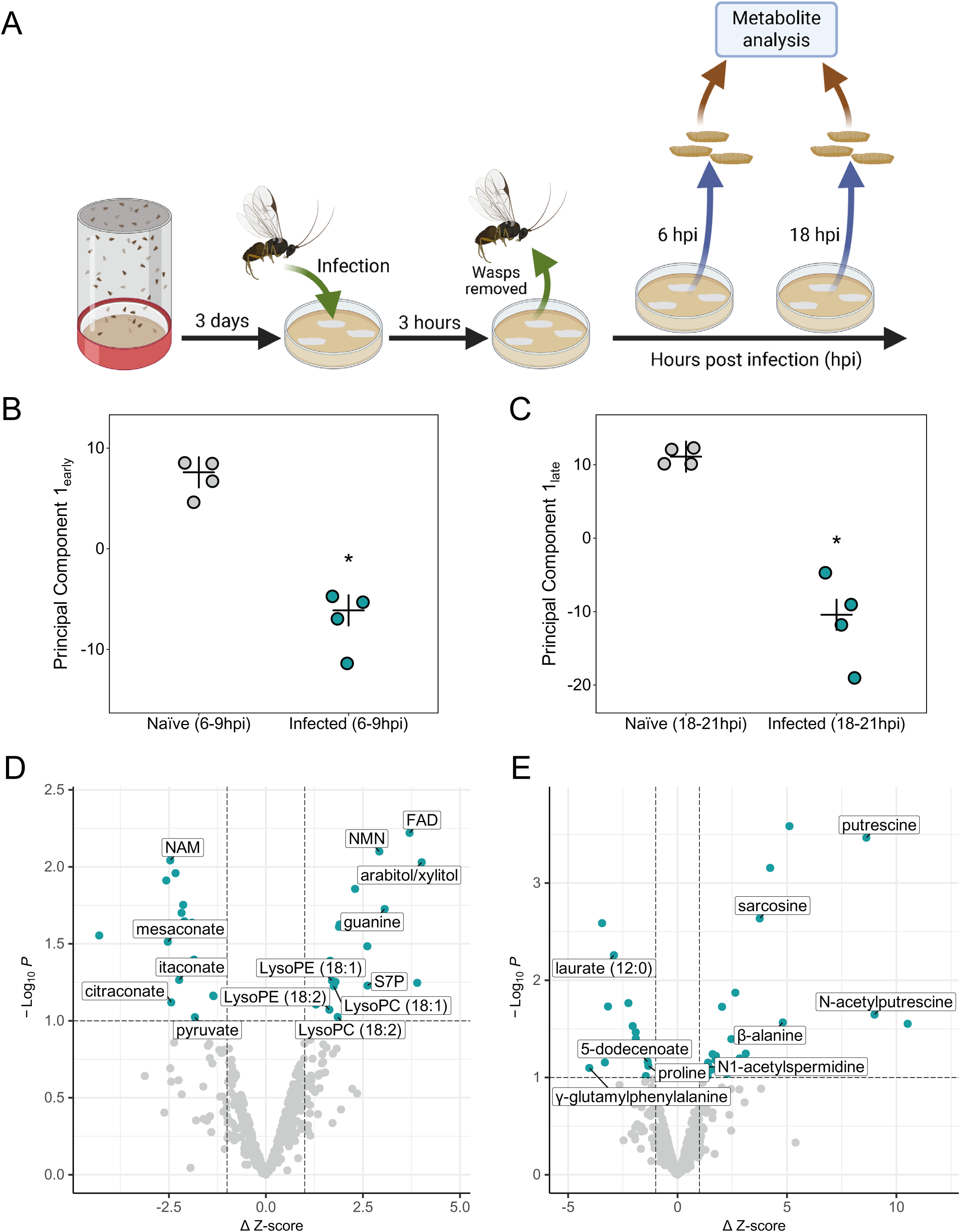
Global metabolomic analysis of immunometabolic signatures following infection. (A) Schematic of sample collection scheme. (B-C) Plots of PC1 values from principal components analysis of metabolites at 6-9hpi (B) and 18-21hpi (C). * indicates p < 0.05. (D-E) Volcano plots of changed metabolites at 6-9hpi (D) and 18-21hpi (E). See also Figures S1-S4.

In total, we were able to identify 447 metabolites from our *D. melanogaster* larval samples (Table S1). Principal components analysis (PCA) of metabolite abundance revealed significant differences in metabolite composition between infected and naïve larvae at both 6-9hpi and 18-21hpi. The first principal component at 6-9hpi (PC1_early_) explained 64% of the variation between the samples with significant differences between naïve and infected larvae (Figure 2B; p = 5.06x10^-4^). Likewise, PC1_late_ explained 76% of the variation and significantly varied between naïve and infected samples at 18-21hpi (Figure 2C; p = 4.14x10^-3^). These PC1 scores are driven by individual metabolites with significant differences in abundance between naïve and infected larvae, and accordingly, we found 68 metabolites that showed a significant change in abundance with infection at either time point (at a significance level of α < 0.1, and ΔZ-score > 1; Table S1). Our global metabolomic data support previous findings focused on specific metabolite classes, namely changes in lipid and glucose metabolism following infection ^4,28,44^, and identify further changes in amino acid and nucleotide metabolism.

We found 36 metabolites that were changed in infected larvae at 6-9hpi, with 17 metabolites showing significantly increased abundance and 19 metabolites showing significantly decreased abundance compared to naïve controls (Figure 2D). Notably, there is evidence for a conserved switch in glucose metabolism away from the energy producing glycolysis: tricarboxylic acid (TCA) cycle and towards the anabolic PPP. Specifically, we observed an increased abundance of the PPP metabolite sedoheptulose-7-phosphate (S7P; Figure S1A), and potential downstream PPP products including arabitol/xylitol (Figure S1B), the nucleotide guanine (Figure S1C), and several lipid species (Figure S2A-D). Our findings agree with previous studies of immunometabolism, with increased S7P levels also observed in the hemocytes of infected *D. melanogaster* and activated human immune cells ^44,53^. Notably these studies used targeted metabolomics and did not report levels of the other glucose metabolites we identified. Conversely, we found a decreased abundance of the glycolysis product pyruvate (Figure S1D), and the TCA cycle derivatives mesaconate (Figure S1E), citraconate (Figure S1F), and itaconate (Figure S1G). This altered glucose metabolism is widely conserved in activated immune cells from mammalian macrophages and neutrophils to *D. melanogaster* hemocytes ^5,44,53^. Previous work in the infection-PPP association in *D. melanogaster* was performed using infection by the virulent parasitoid *Leptopilina boulardi* ^44^, and our results with an avirulent parasitoid species support the conclusion that the reallocation of glucose to PPP is an immunometabolic adaptation rather than a consequence of parasitoid venom activity.

The 6-9hpi time point also displayed altered levels of lysophospholipids (LPLs; Figure 2D), lipid signaling molecules which may play a conserved role in immunity ^54^. We found an increased abundance of several unsaturated 18-carbon LPL species including lysophosphatidylethanolamine (LysoPE; Figure S2A-B) and lysophosphatidylcholine (LysoPC; Figure S2C-D). The production of LPLs is increased by cellular stress, including infection, and LPLs are proposed to act as signaling molecules in regulating inflammatory responses ^54,55^. LPLs bind to specific G protein-coupled receptors (GPCRs) and activate distinct downstream immune and inflammation associated genetic programs ^55,56^. Altered lipid profiles have been previously observed in infected *D. melanogaster*, with infection resulting in a reallocation of lipid anabolism from storage molecules (e.g., triacylglycerols) to the production of phospholipids ^28^. This switch is mediated by activation of the Toll immune signaling pathway, leading to expression of Kennedy pathway enzymes and subsequent production of phospholipids, namely phosphatidylethanolamine (PE) and phosphatidylcholine (PC) ^28^. LysoPE and LysoPC levels were not assessed in this previous work, and our findings suggest that some amount of the PE and PC produced following infection are likely converted to LysoPE and LysoPC via the Lands’ cycle ^57^ and may function as immune signaling metabolites. Interestingly, the expression of LPL-specific GPCRs is also regulated by Toll-like Receptor signaling ^55^. Previous work suggests that the Toll signaling pathway is activated at early time points following parasitoid infection ^22,29^, providing a possible basis for the shift in lipid metabolism following infection.

We observed evidence for activation of the nicotinamide adenine dinucleotide (NAD+) salvage pathway at 6-9hpi. In this pathway, cellular nicotinamide (NAM) is converted to nicotinamide mononucleotide (NMN), the precursor molecule for NAD+ synthesis ^58^. We found a decrease in NAM and concomitant increase in NMN in *L. clavipes* infected larvae at 6-9hpi compared to naïve larvae (Figure S2E-F). NAD+ is required as a cofactor for a multitude of cellular reactions, and the immune induced activation of the salvage pathway suggests that NAD+-mediated processes play an important role in immune activation following parasitoid infection. This reliance on NAD+ is also likely conserved, with a noted role for NAD+ in the mammalian immune response to infection ^59^.

As expected from the PCA results (Figure 2C), we also found significant changes in individual metabolites at the late time point following infection. At 18-21hpi we found 36 metabolites that were significantly altered by infection (Figure 2E); 23 of these metabolites had an increased abundance whereas 13 metabolites were decreased in comparison to naïve controls. We additionally found that 4 of our identified metabolites showed significant changes in abundance at both timepoints (Table S1). Interestingly, at 18-21hpi we see evidence for decreased abundance of energy storage molecules, such as laurate (Figure S3A) and 5-dodecenoate (Figure S3B). This finding is in line with proposed life history trade-offs in which resources are reallocated for immediate use by the immune response rather than storage for subsequent use in development or reproduction.

We additionally found evidence for altered protein and amino acid metabolism at 18-21hpi, including altered levels of sarcosine (Figure S3C), proline (Figure S3D), β-alanine (Figure S3E), and γ-glutamyl amino acids (Figure S3F-G), suggesting alterations in protein stability and amino acid transport. This altered amino acid metabolism is also reflected in the increased abundance of the polyamine putrescine (Figure S4A), its precursor N1-acetylspermidine (Figure S4B) and its by-product N-acetylputrescine (Figure S4C). The increase in polyamines may also represent a conserved immune process; they are important for immune responses in both plants and animals^60,61^.

Finally, we wanted to test whether the immunometabolic changes we observed were associated with larval developmental changes between the 6-9hpi and 18-21hpi time points. We found that 62 metabolites changed abundance with development, with 35 showing greater abundance at 6-9hpi and 27 showing greater abundance at 18-21hpi. 18 of these metabolites are in common with our identified immunometabolites, and interestingly, 11 of these metabolites show an inverse relationship between development and immunity (Figure S5; Table S2). This discordance may further reflect the reallocation of metabolic resources from development to immunity following infection.

### PVF signaling results in the production of an immunometabolic-like state

Because *Pvf1* has been previously linked to a type of metabolic switch in *D. melanogaster* development ^46^ and loss of PVF signaling led to decreased immune competence (Figure 1A-D), we hypothesize that PVF signaling may play a role in metabolic reallocation following infection. If PVF signaling is the immunometabolic switch, we would expect that ectopic activation of PVF signaling would result in larvae with a similar metabolite profile to *L. clavipes* infected larvae. To test this prediction, we performed global metabolomic analysis on *eater>Pvr.λ* larvae at the same developmental time points as in Figure 2A. Constitutive PVF signaling led to significant alterations of 176 metabolites compared to controls (Table S3), including 87 metabolites with significantly altered abundance at the early time point, and 123 metabolites with significantly altered abundance at the late time point. 34 of the *eater>Pvr.λ* responsive metabolites had a significantly altered abundance at both time points. We performed PCA analysis of metabolite abundance to explore the relationship between our samples. We found that the three conditions (naïve, *L. clavipes* infected, and *eater>Pvr.λ*) clustered independently at the early time point (Figure 3A) but that there was overlap between infected and constitutive PVF at the late time point (Figure 3B).

**Figure 3.**
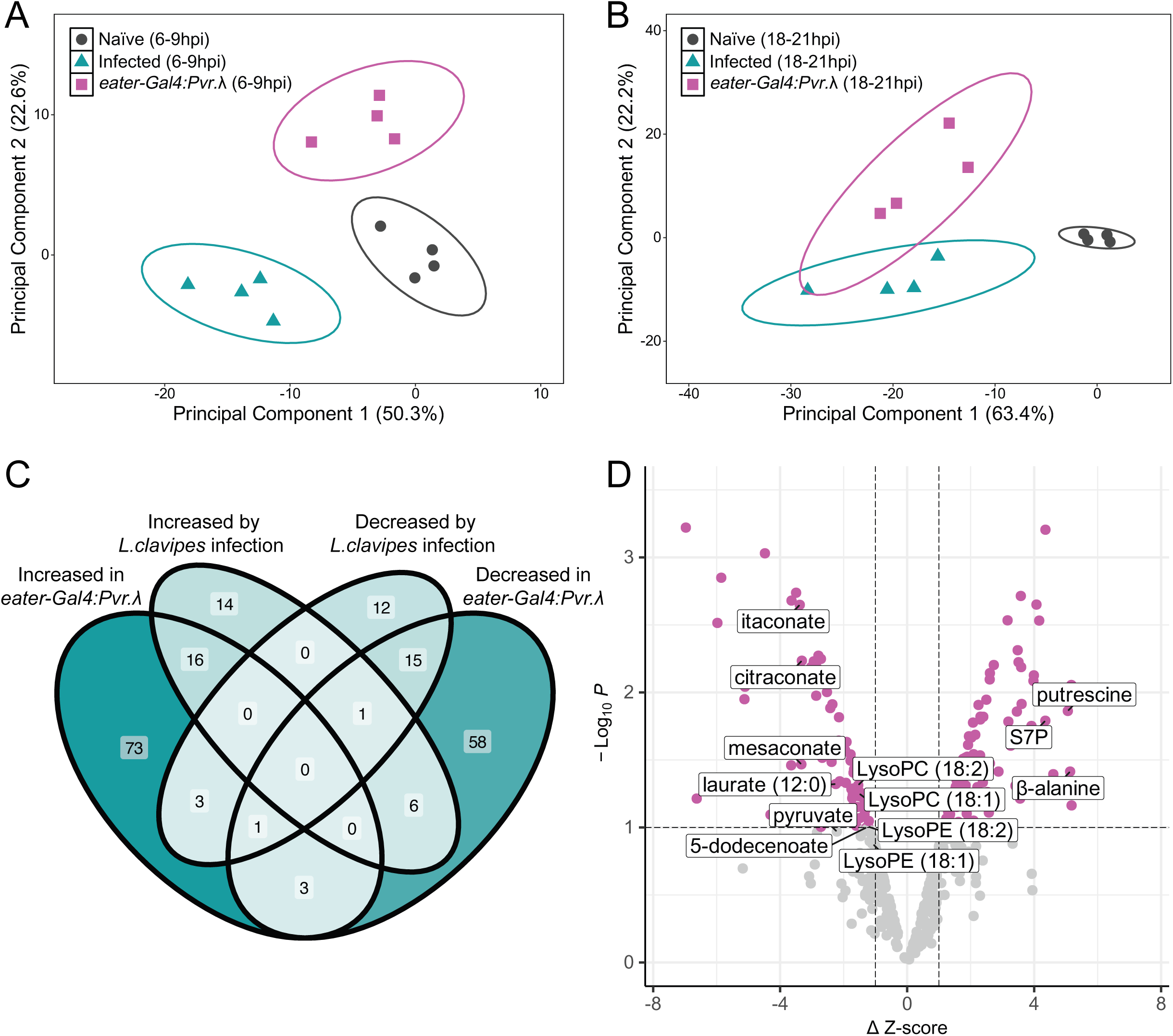
PVF signaling induces an immunometabolic-like state. (A-B) Plots of the first two principal components from principal components analysis of metabolites at 6-9hpi (A) and 18-21hpi (B). (C) Venn diagram comparing metabolite changes in *L. clavipes* infected and *msn>Pvr.λ* larvae. (D) Volcano plot showing the abundances of infection-dependent metabolites in *msn>Pvr.λ* larvae.

As expected from on the PCA results, metabolites showing changed abundance following *L. clavipes* infection were significantly enriched among the metabolites altered in *eater>Pvr.λ* larvae (42 of 68 identified metabolites; p = 4.34x10^-5^). Immunometabolites were enriched among the PVF-responsive metabolites that both increase (p = 1.54x10^-3^) or decrease (p = 5.49x10^-6^) in abundance following infection and *eater>Pvr.λ* expression (Figure 3C). Despite the independence of PCA scores at the early time point, we found an enrichment of immunometabolites among the PVF-responsive metabolites at both the early (p = 0.0292) and late (p = 1.98x10^-4^) time points. The significant overlap between *L. clavipes*-induced and *eater>Pvr.λ*-induced metabolic changes (Figure 3D) suggests that PVF signaling plays an important role in mediating the metabolic reallocation following parasitoid infection.

### Metabolic reallocation mediates an immunity: development trade-off in *D. melanogaster*

In *D. melanogaster* life history, energy stores established during larval development are essential for the completion of pupal development ^62,63^. If the PVF-mediated changes in host metabolism following infection lead to a metabolic trade-off, then we would expect to observe altered life-history traits subsequent to the induction of the immunometabolic state. To test this prediction, we expressed *msn>Pvr.λ* to mimic a chronic immunometabolic state and compared post-larval development to a genetic background control. We also assessed post-larval development in *msn>Pvr.DN* flies to test if PVF signaling is required for development during this stage. We tracked *D. melanogaster* development from the late larval stage (L3) using three developmental milestones: the onset of pupal development (P1), the pharate adult stage (P14), and eclosion of adult flies (Figure 4A) ^64^.

**Figure 4.**
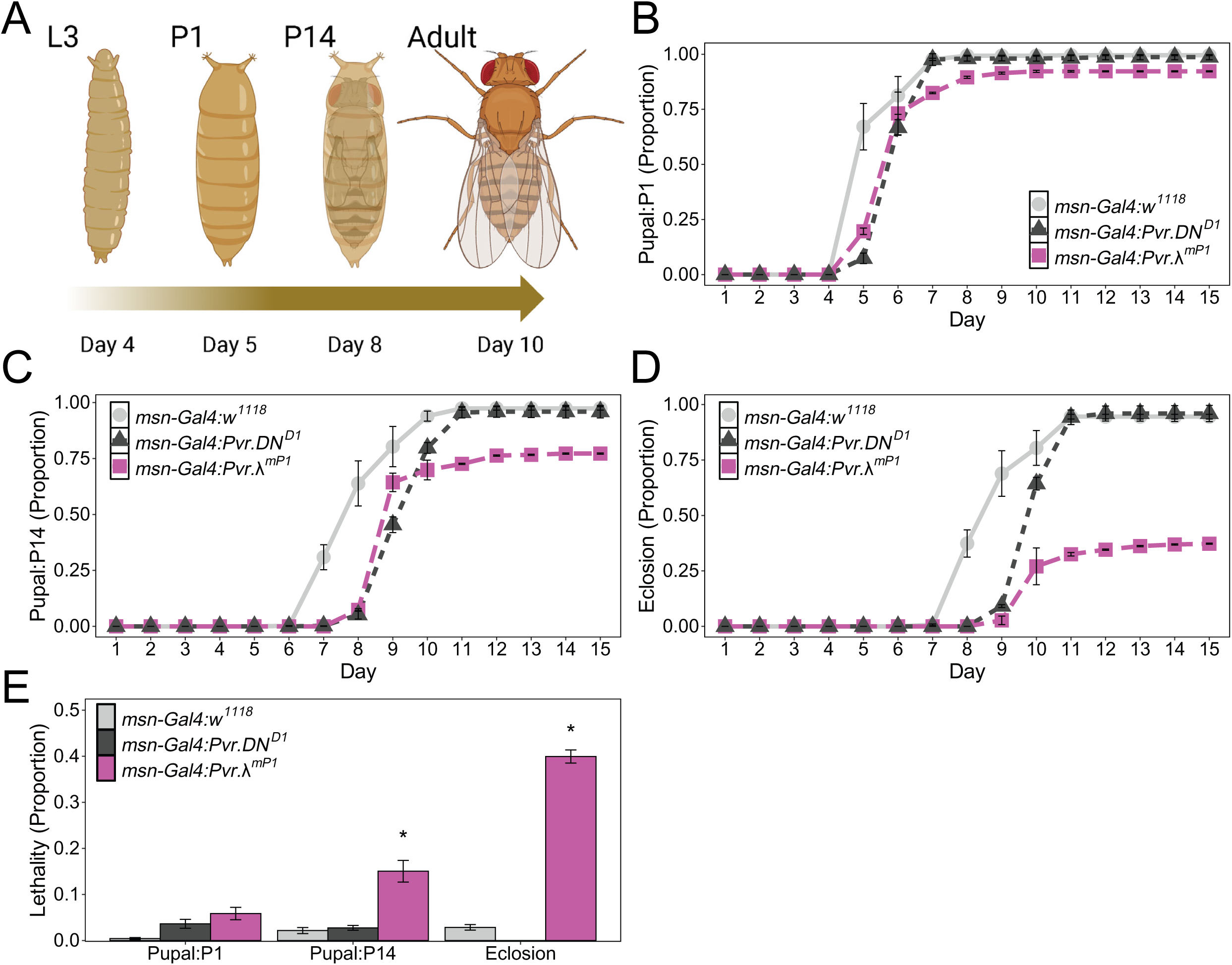
An immunity: development trade-off is mediated by PVF signaling in *D. melanogaster*. (A) Schematic of *D. melanogaster* developmental stage analyzed. (B-D) Plots of cumulative proportions of larvae reaching P1 (B), P14 (C), and adult (D) developmental stages on the indicated day. (E) Plot of cumulative lethality at each developmental stage in the indicated genotypes. * indicates p < 0.05 by Dunnet’s test compared to control.

We found that expression of the constitutively active *msn>Pvr.λ* impaired post-larval development. *msn>Pvr.λ* larvae were significantly delayed in reaching P1 compared to genetic controls (Figure 4B; p = 6.67x10^-6^ on day 5). Interestingly, a similar delay was observed for *msn>Pvr.DN* expressing larvae (Figure 4B; p = 1.10x10^-7^ on day 5). These developmental delays continued throughout and were lengthened when assessing development to the P14 stage. *msn>Pvr.λ* flies were two days slower in reaching P14 compared to genetic controls (Figure 4C; p = 3.42x10^-3^ on day 7, and p = 2.13x10^-7^ on day 8). Again *msn>Pvr.DN* flies showed a similar lengthened delay (Figure 4C; p = 5.77x10^-3^ on day 7, p = 3.10x10^-7^ on day 8, and p = 1.73x10^-3^ on day 9). As expected, flies with manipulated PVF signaling also showed delayed eclosion. *msn>Pvr.λ* flies were two days slower to begin eclosing compared to genetic controls (Figure 4D; p = 9.51x10^-4^ on day 8, and p = 1.66x10^-8^ on day 9). *msn>Pvr.DN* flies showed a similar delay (Figure 4D; p = 1.79x10^-3^ on day 8, and p = 9.55x10^-7^ on day 9). These results suggest that PVF signaling must be finely tuned for *D. melanogaster* post-larval development, and that either reduced or ectopic PVF activity can result in a developmental delay.

Interestingly, beyond the delay, we observed that *msn>Pvr.λ* flies failed to reach P14 and eclose in comparable numbers to the genetic controls (Figure 4C-E). We found that only 85.0 +/-2.4% of *msn>Pvr.λ* expressing flies reached P14, which reflected significant lethality compared to the genetic control flies with 97.8 +/-0.07% reaching P14 (Figure 4E; p = 0.0311). A stronger degree of lethality was observed at eclosion, with only 60.1 +/-1.4% of *msn>Pvr.λ* flies reaching adulthood, compared to 97.1 +/-0.06% of controls (Figure 4E; p < 1x10^-10^). In contrast, *msn>Pvr.DN* expression did not cause significant lethality at any developmental stage assayed (Figure 4E). These findings support a role for PVF signaling in an immunity: development trade-off, by establishing that flies experiencing an immunometabolic state demonstrate decreased survival to adulthood and delayed developmental rate.

### Survival of virulent parasitoid wasps is independent of host PVF signaling

Our results suggest that PVF pathway activity is necessary for anti-parasitoid immunity and that ectopic PVF signaling is sufficient to induce an immunometabolic state in *D. melanogaster* larvae. We next wanted to test whether this state would confer an advantage to larvae when confronted with virulent wasps. If host immunometabolic state influenced their ability to survive infection, we would predict that *msn>Pvr.λ* expression would lead to decreased wasp eclosion from infected fly larvae. Alternatively, parasitoids are dependent on their host for nutrition, and so Drosophila-infecting parasitoids may have evolved to depend on the PVF-mediated immunometabolic state. If virulent parasitoids require host immunometabolism for survival, we predict that *msn>Pvr.DN* expression would lead to decreased wasp survival and eclosion compared to the genetic background control.

To test these competing ideas, we infected *msn>Pvr.λ* and *msn>Pvr.DN* larvae with the virulent parasitoid species *Leptopilina heterotoma* (Figure 5A) and *Ganaspis hookeri* (Figure 5B), and assessed wasp eclosion compared to the genetic background control. We observed that both wasp species emerged from *msn>Pvr.λ* and genetic control flies at a comparable rate (Figure 5A-B). This suggests that ectopic PVF signaling alone does not provide a protective effect to larvae encountering virulent parasitoids. Accordingly, we observed that levels of melanization, a measure of immune activity, did not differ between the genotypes (Figure 5C).

**Figure 5.**
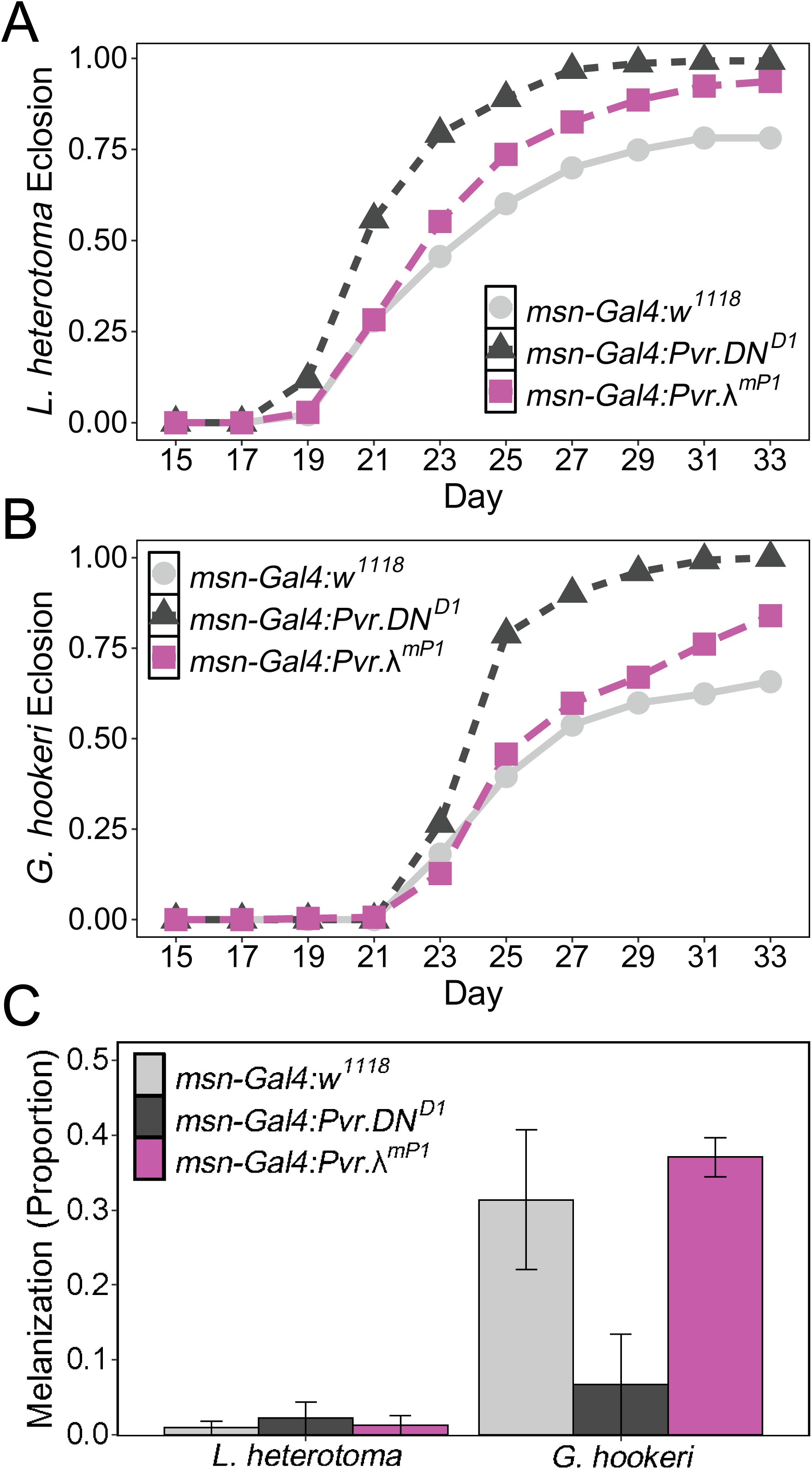
Parasitoid wasp survival is not responsive to host PVF signaling. (A-B) Cumulative proportion plots of parasitoid eclosion (A: *L. heterotoma*; B: *G. hookeri*) on the indicated day. (C) Proportion of melanized larvae infected by the indicated parasitoid species.

We further found that both wasp species were able to eclose from *msn>Pvr.DN* flies with no identifiable decrease in parasitization success (Figure 5A-B). This finding suggests that rather than relying on host immunometabolism, parasitoids may instead have evolved to manipulate host metabolism. This idea is supported by the discovery of parasitoid venom proteins with homology to metabolic enzymes ^65–68^. Interestingly, immune activity, as assessed by melanization rate, was decreased in *msn>Pvr.DN* flies (Figure 5C), in agreement with previous findings that the melanotic encapsulation response was impaired in this genotype (Figure 1B).

## DISCUSSION

### Immunometabolism and the response to parasitoid infection

Our results reveal that parasitoid-infected *D. melanogaster* undergo a general metabolic reallocation, leading to altered levels of a wide variety of metabolites. This metabolic switch is primarily initiated by release of Pvf1 from immune-activated hemocytes (i.e. *msn*-expressing hemocytes; Figure 1C-D), and is mediated by Pvr activation in both circulating plasmatocytes and immune activated hemocytes (Figure 1A-B). This dependence on immune activation for triggering immunometabolism has the effect of coupling the metabolic alterations to infection, and may help to prevent inappropriate triggering of the immunometabolic state. We further found that a successful immune defense requires the switch to immunometabolism, in agreement with previous findings that the activation of immune cell responses across species is at least partially dependent on metabolic changes ^8^. This altered metabolism is associated with shunting resources to immune cells; in naïve *D. melanogaster* larvae ∼10% of carbon derived from dietary glucose intake is incorporated into hemocytes, but this increases to ∼30% in infected larvae ^4^. Our findings support a model in which this spatial reallocation is accompanied by changes to the metabolite pool itself, likely via differential use of metabolic pathways including the activation of PPP in glucose metabolism and the Kennedy pathway and Lands’ cycle in lipid synthesis (Figure 2) ^28,44^.

Our global metabolomic analysis has allowed us to identify more than 60 metabolites whose abundance changes with infection status. These metabolites display a high degree of overlap with previously identified immunometabolism changes in glucose and lipid metabolism, NAD+ salvage, and polyamine accumulation. However, there are some limitations to this approach which warrant discussion. A limitation of metabolomic studies in general, is that for any approach not all metabolites are able to be detected, and many of the detected compounds cannot be identified as specific metabolites ^69^. Global metabolomics additionally may not have sufficient sensitivity to detect or quantify lower abundance metabolites, but can be used as an unbiased approach to detect metabolite changes, in contrast to the limited scope but greater sensitivity provided by targeted metabolomics ^70,71^. The resulting data need to be interpreted carefully, and can be extrapolated to provide insight into the affected pathways, but may not give an in-depth view of every metabolite change.

A majority of the identified immunometabolites were also found to be altered in larvae with constitutive PVF signaling (Figure 3), and these likely represent the metabolic changes that are dependent on the PVF-mediated metabolic switch. However, it is also interesting to consider those immunometabolites which appear to be regulated independent of PVF signaling. They may reflect metabolites that are regulated via a distinct immune mechanism, or that change as part of a parasitoid species-specific response. Notably, the LPL species that are elevated in *L. clavipes* infected larvae (LysoPE and LysoPC) show a decreased abundance in larvae with constitutive PVF signaling (Figures 2D and 3D). Activation of the Kennedy pathway, and subsequent changes in lipid synthesis have been linked to Toll pathway activity ^28^, suggesting that other cytokines and immune pathways contribute to establishing the immunometabolic state.

It is also likely that not every metabolite identified in a metabolomic screen is directly involved in the process examined. For instance, identified metabolites may accumulate as a by-product of an upstream metabolic process that is directly involved. In the case of immunometabolism, the altered metabolites may also act through various mechanisms, including the transfer of energy, immune cell signaling, or even as an immune effector, or they may be involved in the post-resolution recovery from the immune state rather than the defense response itself. Thus it will be important to test the roles of identified metabolites of interest. This reveals a strength of our experimental system, with *D. melanogaster* providing an excellent genetic model to test manipulations to metabolic pathways. As an example, we tested the role of polyamines in antiparasitoid immunity by infecting *Ornithine decarboxylase 1* (*Odc1*) mutant larvae. The *Odc1^M^*^I10996^ mutation has previously been shown to block polyamine synthesis ^72^, and yet has no effect on larval encapsulation ability following *L. clavipes* infection (p = 1.0; 55% encapsulation in *Odc1^MI^*^10996^ vs 58% in the *y,w* genetic background control). While this result requires additional study to further characterize the role of polyamines in immunity, it is possible that polyamines are redundant with another immune or immunometabolic mechanism, or that polyamines play an accessory role in determining fitness or life history trade-offs, rather than acting in the encapsulation response. Polyamine levels are tightly regulated throughout development ^73^, and so the infection induced polyamines may instead be important for continued host development after the resolution of the infection. Finally, the polyamines that accumulate in hosts following infection may be synthesized via a distinct pathway ^73^ and not suppressed in our *Odc1* mutant.

Similarly, parasitoids or other pathogens may evolve to tolerate host immunometabolic mechanisms. The virulent *Drosophila*-infecting parasitoid wasps have highly evolved to successfully infect hosts, using venom proteins to block host immune mechanisms at various stages ^31,74–77^. It therefore may not be surprising that constitutive PVF signaling is not sufficient to confer host protection. It does suggest that if flies are using metabolites as immune effectors or toxic molecules, the wasps have evolved to detoxify or otherwise inhibit this effect of the metabolite. It further suggests that if flies use immunometabolism to deprive developing parasitoids of essential nutrients, the wasps have evolved to counteract this reallocation and ensure access to the necessary metabolites. This mechanism may also be reflected in the lack of impact of dominant negative PVF signaling on wasp survival. This finding suggests that developing wasps are not dependent on their hosts’ metabolic allocation, as has been observed for other pathogen species ^78^. These observations suggest that parasitoid wasps may have evolved to manipulate host metabolism to promote successful parasitization. This is supported by the multi-omic profiling of *G. hookeri* and *L. heterotoma* venoms, which has identified venom proteins showing strong conservation with known metabolic enzymes, and functional analysis of the Leptopilina-specific Venom Lipase protein which acts to hydrolyze host lipids in order to provide required nutrients for wasp embryogenesis ^31,66–68^. Interestingly, both of the virulent species we tested in our analysis of PVF signaling, *G. hookeri* and *L. heterotoma*, are generalist parasitoid species and able to infect a wide range of *Drosophila* species. It would be interesting to test whether phylogenetically restricted specialist parasitoid species would have an elevated sensitivity to or dependence on host metabolic state.

### *D. melanogaster* life-history and the immunity: development trade-off

The life cycle of holometabolous insects, including *D. melanogaster*, leads to a unique life-history in which different developmental stages may have distinct nutritional requirements, environments, and even opportunities to feed. During the larval stage, *D. melanogaster* must support growth and proliferation for development, and accumulate energy stores for future post-larval development ^62^. As a consequence, larval metabolism functions to convert acquired nutrients into anabolic building blocks while also shunting considerable nutritional resources into synthesis of energy storage molecules including glycogen and triacylglycerides (TAGs) ^62^. Interestingly, larvae are able to continue to develop during periods of nutrient stress by mobilizing this stored energy, however larvae raised on nutrient poor diets have an increased developmental time ^62,63,79^. It has been hypothesized that chronic activation of immunometabolism would lead to nutrient deprivation and subsequent pathogenesis of the host ^8^. The high levels of lethality resulting from constitutive PVF signaling (Figure 4E) support this hypothesis.

The first milestone of post-larval development is the initiation of pupation (P1 stage). Trehalose is needed to promote pupation, and the onset of pupation is dependent on levels of trehalose, rather than storage molecules such as glycogen ^62^. The pupation delay seen in infected or constitutively active Pvr expressing larvae (Figure 4B-D) is therefore likely due to the glucose metabolism changes seen following infection (Figure 2) ^4,43,44^. Following pupation, the metabolic rate of the developing fly drops to a minimal level and stays low, until increasing again as the fly nears the end of metamorphosis (i.e., approaches eclosion) ^80^. At this stage, the pupa largely relies on the mobilization of lipid stores for energy ^62,80^. Because pupal development is a non-feeding stage, lethality during the final pupal stages or at eclosion is likely due to insufficient lipid storage during the larval stage. The activation of the Kennedy pathway in response to infection would shift lipid metabolism away from TAG synthesis, potentially explaining the P14/eclosion stage lethality we observed in constitutively active Pvr expressing flies (Figure 4E).

The observed relationship between the immune induced metabolic reallocation and organismal development supports the existence of an immunity: development trade-off in *D. melanogaster*. In our experiments, the expression of constitutively active Pvr mirrors the effect of infection, but represents a chronic switch to the immunometabolic state, and therefore appears to be more extreme than the temporary reallocation mediated by infection with additional metabolic changes, larger effect sizes, and lack of a post-resolution recovery period. This chronic immunometabolic condition may result in a more exaggerated trade-off phenotype. In agreement with this, flies mounting a successful immune defense against parasitoid infection survive to adulthood, but do show life-history trade-offs; as adults they have a smaller body size and reduced fecundity ^27^. Additionally, artificial selection for flies with better immune success against parasitoid infection show trade-off with reduced larval competitive ability on nutrient limited food ^39,40^. To better understand the impact of metabolic reallocation on life-history trade-offs in the absence of pathogen infection, the expression of constitutively active or dominant negative Pvr can be finely tuned temporarily using a conditional bipartite expression system in *D. melanogaster* ^81^. This would allow for the study of a more ecologically relevant response duration to untangle the specific effects of metabolic reallocation from other effects that pathogen infection may have on life-history trade-offs.

### The genetic basis of immune-mediated metabolic reallocation: A conserved role for Pvf signaling?

Our study has revealed a high degree of conservation of immunometabolic changes among host species, and may highlight a possible conserved role for PDGF/VEGF/Pvf signaling in regulating organismal metabolism in response to environmental cues. In *D. melanogaster*, the PVF signaling pathway has been linked to metabolic regulation in response to developmental signaling ^46^, dietary conditions ^47^, tumorigenesis ^48^, and infection. In these mechanisms, PVF cytokines are released in response to a stimulus and act to alter the balance of nutrient storage. This metabolic regulatory role appears conserved across species, with both of the derived PDGF and VEGF signaling pathways regulating with lipid accumulation in mammalian models ^47,82^. Interestingly, PDGF/VEGF/Pvf signaling is able to either promote or prevent the accumulation of lipid stores, with the context determining the outcome. For instance, in *D. melanogaster*, the *Pvf1* ligand is associated with decreased energy stores following developmental ^46^ and infection cues, whereas the *Pvf3* ligand promotes TAG accumulation in response to dietary stress ^47^. The identity of the initiating and target tissues, and mode of signaling may also be determinants of PVF signal outcome.

The role of PVF signaling in immunometabolism appears to be unique in having an autocrine/paracrine mode of signaling. Despite this putative restriction of PVF signaling to hemocytes, it is likely that the changes we observed are the result of global metabolic changes, potentially involving disparate tissues such as the fat body, gut, and body wall muscle along with the hemocytes. Previous studies have demonstrated that extracellular adenosine (eAdo) can function as an endocrine signal for metabolic reallocation in parasitoid infected *D. melanogaster*^4^. eAdo is released from hemocytes via the ENT2 transporter and signals via the Adenosine receptor (AdoR) ^4^. Although the tissue specificity of this mechanism is thus far unknown, it has been proposed that AdoR expression in the central nervous system or endocrine glands is a central regulator of metabolism ^4^. While a putative interaction between PVF and eAdo remains to be determined, manipulation of eAdo signaling phenocopies alterations in the PVF pathway, with loss of AdoR leading to decreased encapsulation ability ^4^. It is a possibility that PVF activity is upstream of eAdo, with pathway activity resulting in elevated expression of the ENT2 channel. Alternatively, PVF and eAdo could comprise parallel pathways that act in the immunometabolic switch. Similarly, further research will also be required to understand the downstream effect of PVF signaling in energy storage tissues. Previous findings have uncovered a role in immunometabolic reallocation for the Mef2 transcription factor in the fat body ^83^, and the JAK-STAT signal transduction pathway in muscle ^34,43^, suggesting that these mechanisms may function downstream of PVF signaling.

We are also yet to determine the molecules that initiate *Pvf1* expression in hemocytes following infection. Previous studies suggest that damage-associated molecular patterns (DAMPs) are often associated with the activation of immune responses, including metabolic shifts ^7^. DAMPs are host molecules that are only released following tissue perturbation, and can also be defined as endogenous metabolites released from damaged or dying cells ^54,55,84^. It is therefore possible that a metabolite our studies has identified can act as a DAMP in the antiparasitoid response.

## METHODS

### Insect husbandry

Flies were maintained on standard Drosophila medium of cornmeal, yeast, and molasses. The *w*^1118^ (RRID: BDSC_5905), *eater*-*Gal4* (RRID: BDSC_36321), *UAS-Pvr.DN^D1^*(RRID: BDSC_58430), *UAS-Pvr.λ^mp1^* (RRID: BDSC_58428), *UAS-GAL4^RNAi^*(*P{VALIUM20-GAL4.1}attP2*; RRID: BDSC_35784), *UAS-Pvf1^RNAi^*(*P{TRiP.HMS01958}attP40*; RRID: BDSC_39038), *y^1^,w^1^* (RRID: BDSC_1495), and *Odc1^MI109^*^96^ (RRID: BDSC_56103) *D. melanogaster* strains were obtained from the Bloomington Drosophila Stock Center (Bloomington, IN; RRID: SCR_006457). *msn*-*Gal4* was kindly provided by Dr. Robert Schulz.

Parasitoid wasp species *Leptopilina clavipes* (strain LcNet), *Leptopilina heterotoma* (strain Lh14), and *Ganaspis hookeri* (strain GhFl) were used in this study. *L. clavipes* were maintained on *Drosophila virilis* (NDSC_15010–1051.87) hosts, *L. heterotoma* were maintained on *D. melanogaster* (*Canton S*; RRID: BDSC_64349) hosts, and *G. hookeri* were maintained on *Drosophila yakuba* (Kyorin_k-s03) hosts.

### Immune success assay

To assay immune success, parasitic wasp infections were performed as described ^30^. Three adult female wasps were allowed to infect 30 late second instar larvae for a 72-hour period at 25°C in a 35-mm Petri dish filled with *Drosophila* medium. After 72 hours, the larvae were dissected and scored for successful encapsulation of wasp eggs. All infections were performed in triplicate. To measure melanization activity, larvae were exposed to six female parasitoid wasps for a 4-hour infection period at 25°C in a 35-mm Petri dish filled with *Drosophila* medium. After 72 hours, the larvae were scored for the appearance of melanization.

### Development timeline and lethality

Larvae were collected at the third instar stage and transferred to *Drosophila* vials at 25°C to track development. Each vial was imaged every 24 hours, and the number of larvae, pupae, and adult flies were counted. Developmental analyses were performed with naïve flies and data were collected from 5 replicates per genotype.

### Wasp eclosion assay

Flies were allowed to lay eggs in standard *Drosophila* vials for 24 hours at 25°C. After 72 hours of development to the late second instar stage, larvae were exposed to six adult female wasps (and four males in the case of *L. heterotoma*; *G. hookeri* are parthenogenetic). Eclosion of flies and wasps was recorded daily beginning 4 days post-infection and continuing until no further wasp emergence was observed.

### Metabolomics sample preparation

Groups of *w^11^*^18^ larvae were collected at the late second instar stage and infected with *L. clavipes* for 3 hours at 25°C in a 35-mm Petri dish filled with *Drosophila* medium. Wasps removed at end of the 3-hour infection period. Larvae were collected at 6h or 18h post infection period and flash frozen in liquid nitrogen. Naïve larvae were subjected to the same handling and media without wasp exposure. Each biological replicate was composed of 100 larvae, and 4 biological replicates were subjected to further preparation and global metabolomic analysis at Metabolon.

Samples were prepared using the automated MicroLab STAR® system from Hamilton Company. Following homogenization, proteins were precipitated with methanol under vigorous shaking for 2 min (Glen Mills GenoGrinder 2000) followed by centrifugation to remove protein, dissociate small molecules bound to protein or trapped in the precipitated protein matrix, and to recover chemically diverse metabolites. The resulting extract was divided into four fractions: two for analysis by two separate reverse phase (RP)/UPLC-MS/MS methods with positive ion mode electrospray ionization (ESI), one for analysis by RP/UPLC-MS/MS with negative ion mode ESI, and one for analysis by HILIC/UPLC-MS/MS with negative ion mode ESI. Samples were placed briefly on a TurboVap® (Zymark) to remove the organic solvent. The sample extracts were stored overnight under nitrogen before preparation for analysis.

### Metabolite analysis

Samples were analyzed using ultrahigh performance liquid chromatography-tandem mass spectroscopy (UPLC-MS/MS) at Metabolon. All methods utilized a Waters ACQUITY ultra-performance liquid chromatography (UPLC) and a Thermo Scientific Q-Exactive high resolution/ accurate mass spectrometer interfaced with a heated electrospray ionization source and Orbitrap mass analyzer operated at 35,000 mass resolution. The sample extract was dried then reconstituted in solvents compatible to each of the four methods. Each reconstitution solvent contained a series of standards at fixed concentrations to ensure injection and chromatographic consistency. One aliquot was analyzed using acidic positive ion conditions, chromatographically optimized for more hydrophilic compounds. In this method, the extract was gradient eluted from a C18 column (Waters UPLC BEH C18-2.1x100 mm, 1.7 µm) using water and methanol, containing 0.05% perfluoropentanoic acid (PFPA) and 0.1% formic acid (FA). Another aliquot was also analyzed using acidic positive ion conditions, however it was chromatographically optimized for more hydrophobic compounds. In this method, the extract was gradient eluted from the same afore mentioned C18 column using methanol, acetonitrile, water, 0.05% PFPA and 0.01% FA and was operated at an overall higher organic content. Another aliquot was analyzed using basic negative ion optimized conditions using a separate dedicated C18 column. The basic extracts were gradient eluted from the column using methanol and water, however with 6.5mM Ammonium Bicarbonate at pH 8. The fourth aliquot was analyzed via negative ionization following elution from a HILIC column (Waters UPLC BEH Amide 2.1x150 mm, 1.7 µm) using a gradient consisting of water and acetonitrile with 10mM ammonium formate, pH 10.8. The scan range varied slighted between methods but covered 70-1000 m/z.

Raw data were extracted, peak-identified and QC processed using Metabolon’s hardware and software. Compounds were identified by comparison to library entries of purified standards or recurrent unknown entities. The library is based on authenticated standards that contains the retention time/index (RI), mass to charge ratio (m/z), and chromatographic data (including MS/MS spectral data) on all molecules present in the library. Furthermore, biochemical identifications are based on three criteria: retention index within a narrow RI window of the proposed identification, accurate mass match to the library +/-10 ppm, and the MS/MS forward and reverse scores between the experimental data and authentic standards. The MS/MS scores are based on a comparison of the ions present in the experimental spectrum to the ions present in the library spectrum.

### Statistical analysis

All statistical analysis was performed in R ^85^ as follows:

*Plots*: Plots were generated using the *ggplot2*, *ggfortify,* and *ggpubr* R packages ^86–88^. Volcano plots were generated using the *EnhancedVolcano* R package ^89^. Venn diagrams were generated using the *ggVennDiagram* R package ^90^.

*Immune success*: Immune success data were recorded as proportion of wasp eggs encapsulated and control and experimental genotypes were compared by generalized linear models with a quasibinomial distribution. *Post hoc* pairwise comparisons to the control genotype were made using Dunnett’s test in the *multcomp* R package ^91^.

*Lethality and developmental rate*: Developmental rate data were recorded as a cumulative proportion of all individuals reaching the given developmental stage at each 24-hour assessment. Control and experimental genotypes were compared by two-way generalized linear models by assessing the interaction term. Developmental stage lethality data were recorded as proportion of each stage alive and control and experimental genotypes were compared by generalized linear models with a quasibinomial distribution. *Post hoc* pairwise comparisons to the control genotype were made using Dunnett’s test in the *multcomp* R package ^91^.

*Metabolomics data processing and normalization*: Identified metabolites were quantified using the area-under-the-curve of the associated peak. Metabolites detected in < 80% of the samples were removed from further analysis. Missing values were then imputed using 1/5 of the minimum value of the given metabolite ^92^. The resulting data were then normalized using probabilistic quotient normalization (PQN) ^93^ using the *Rcpm* R package ^94^ and tested for normality using the *nortest* R package ^95^. Log transformation was then applied to any non-normal metabolite data. Normalized data were then subjected to auto-scaling (by Z-score) ^92^.

*Metabolomics PCA*: Principal components analysis of metabolite data was performed using the *factoextra* R package ^96^. The values for PC1early and PC1late between naïve and infected larvae were compared using the Welch Two Sample t-test.

*Metabolite abundance*: Metabolism abundance data were compared between the contrasts indicated in the text using the Welch Two Sample t-test. ΔZ scores were calculated as the difference between the average Z-scores of the groups being compared. Degree of overlap (enrichment) between samples was assessed using the hypergeometric test in R.

## Data availability

The data that underlying this study are publicly available from Open Science Framework at https://osf.io/cpwqh/.

## ACKNOWLEDGEMENTS

This work was supported by NIH grant R35 GM133760 (NTM). ALW-S was supported by a Weigel grant from the Phi Sigma Biological Honor Society at Illinois State University. *Drosophila melanogaster* lines obtained from the Bloomington Drosophila Stock Center (NIH P40OD018537) were used in this study. We would like to thank Carrie Marean-Reardon for technical assistance and members of the Venom Biochemistry & Molecular Biology Lab for feedback on the project.

## Notes

https://osf.io/cpwqh/

### Competing Interest Statement

The authors have declared no competing interest.

