## Supplementary figures and images for "The *Drosophila* PDGF/VEGF signaling pathway regulates host immunometabolism in response to parasitoid infection"

### Figure S1

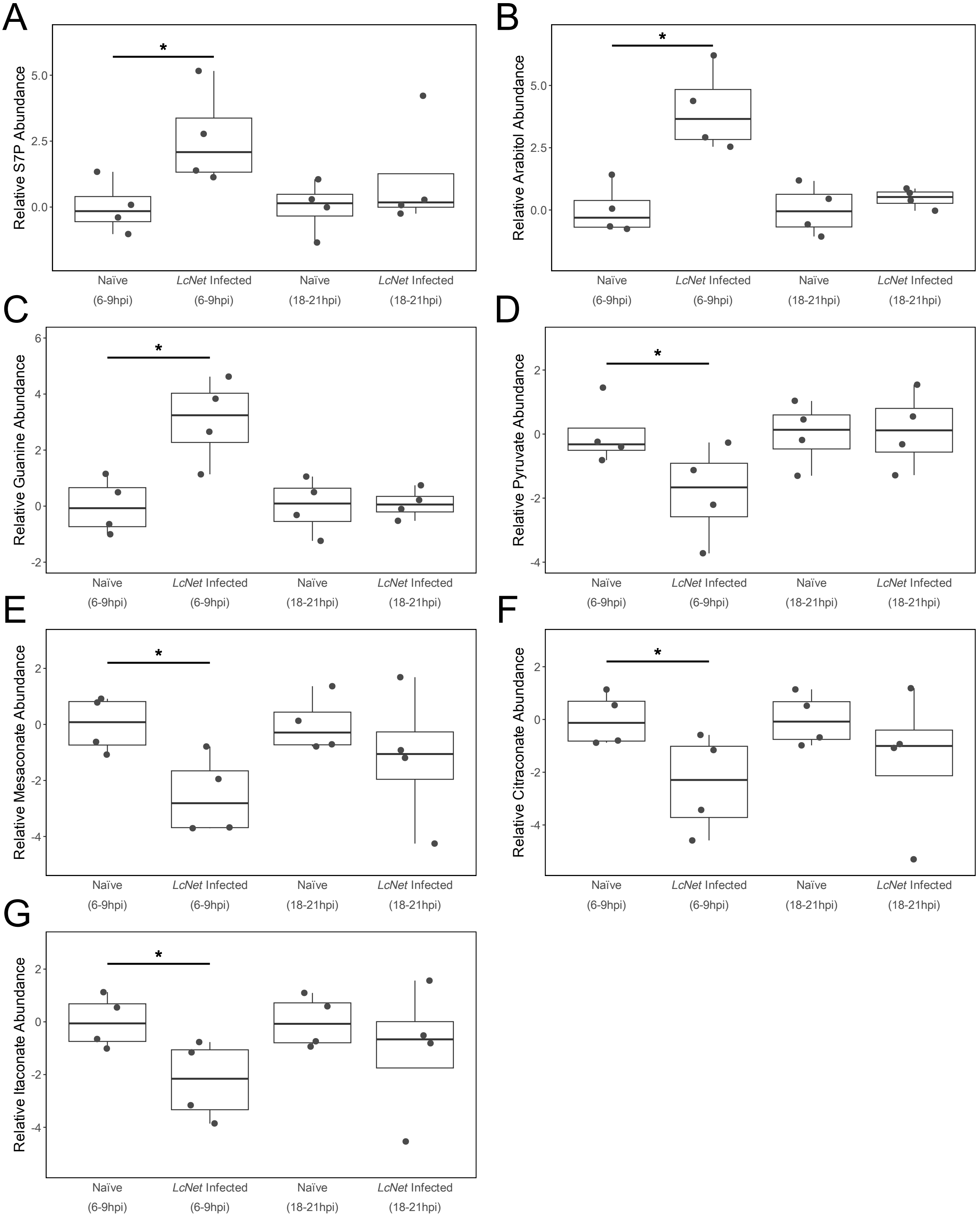

### Figure S2

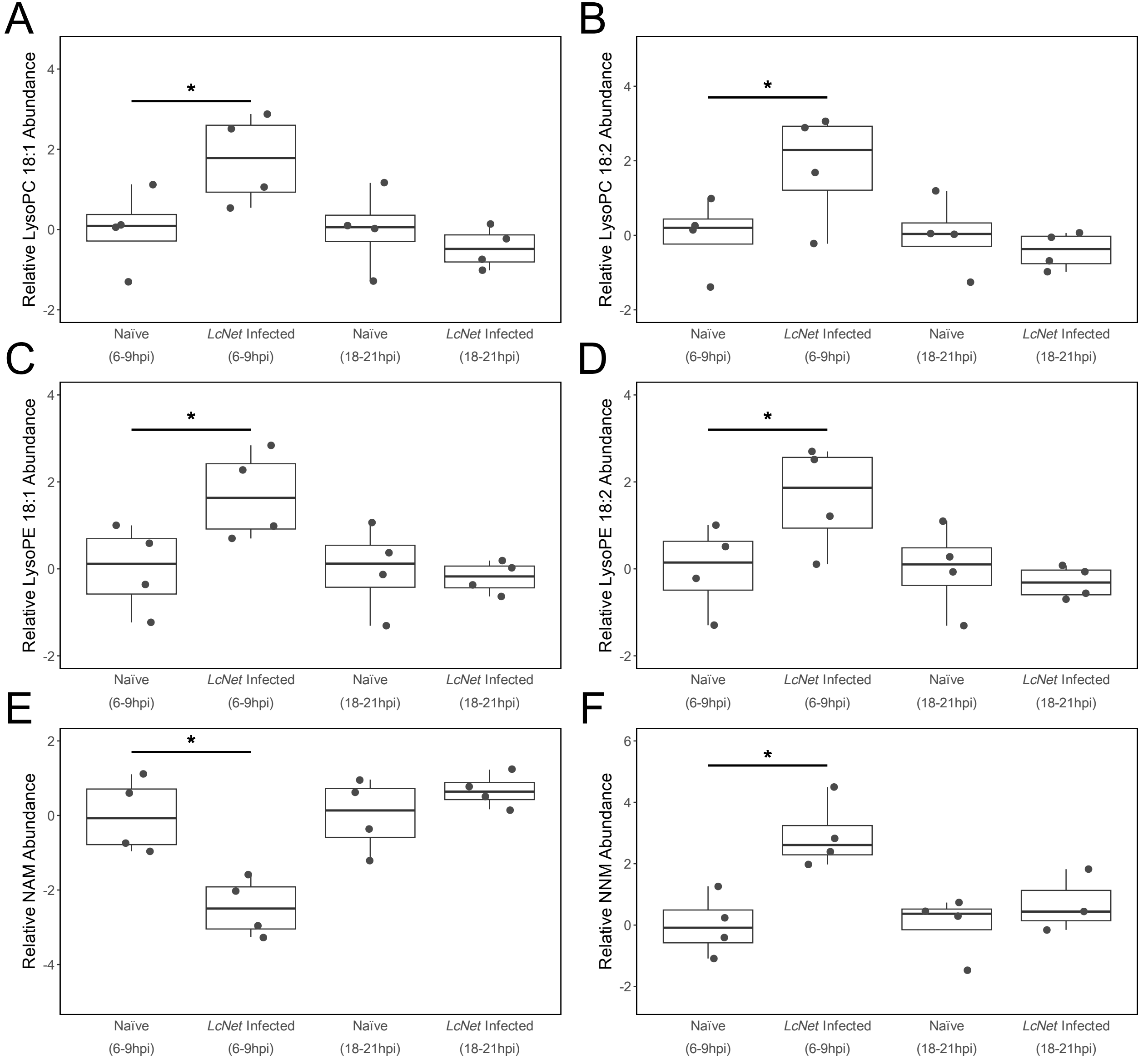

### Figure S3

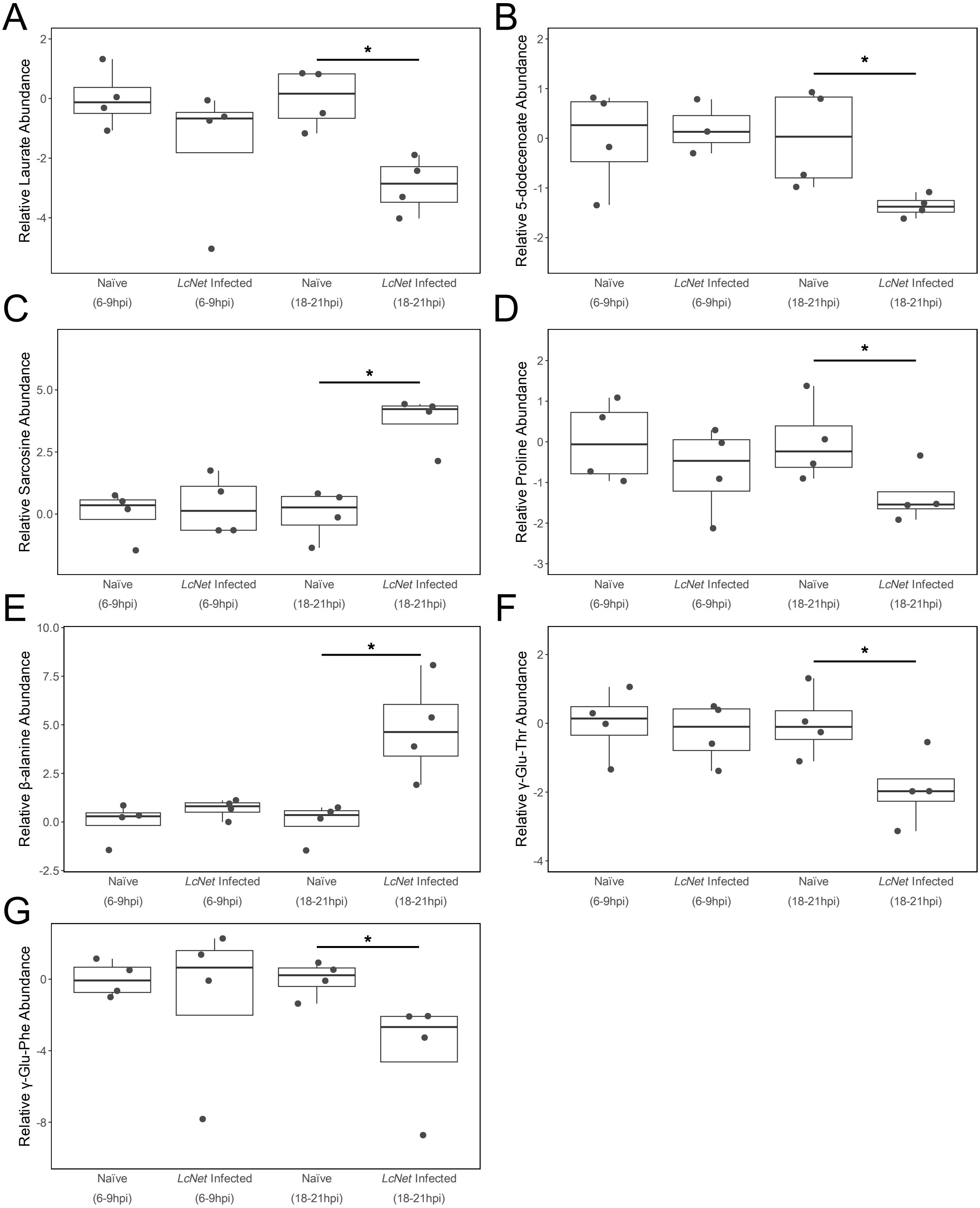

### Figure S4

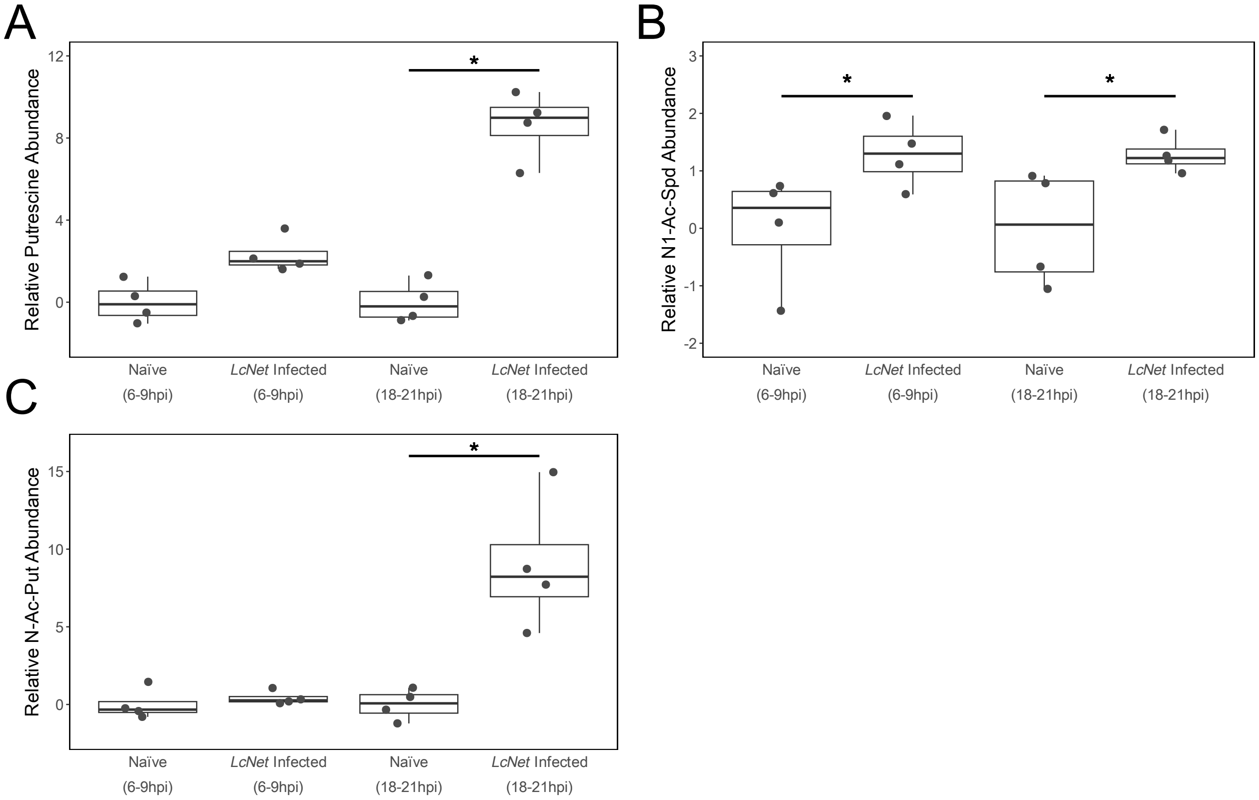

### Figure S5

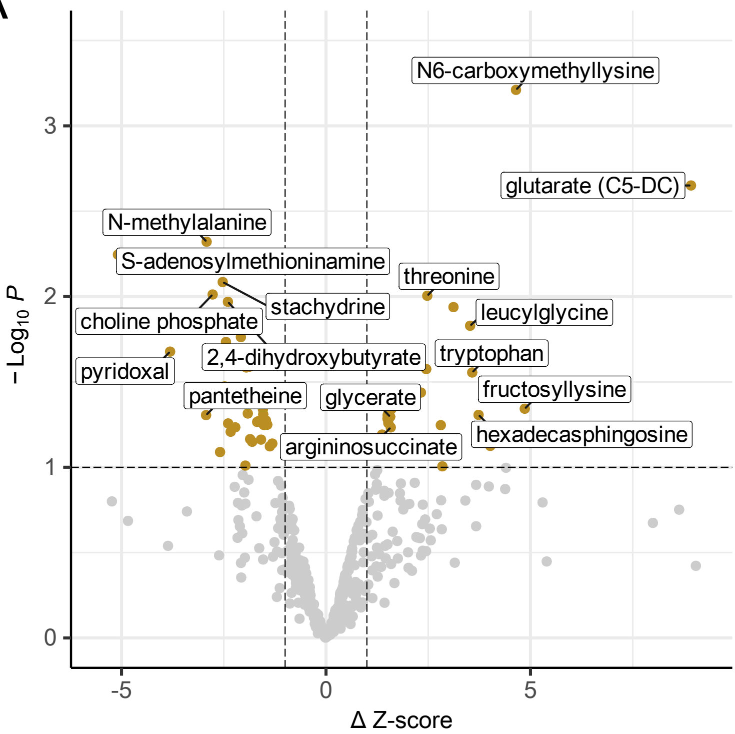

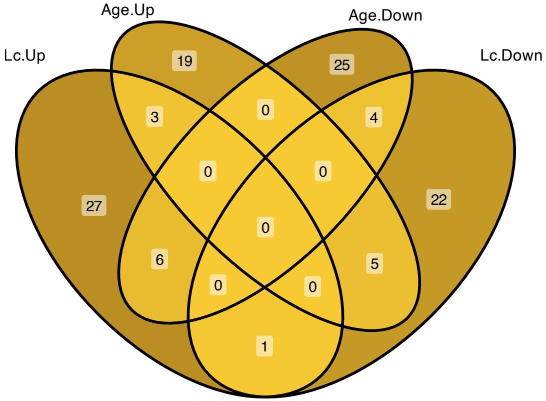
