## Supplemental Information for "The *Drosophila* PDGF/VEGF signaling pathway regulates host immunometabolism in response to parasitoid infection"

**SUPPLEMENTAL FIGURE LEGENDS**

Figure S1. Plots for metabolites involved in glucose metabolism in the indicated samples. (A) S7P, (B) arabitol, (C) guanine, (D) pyruvate, (E) mesaconate, (F) citraconate, and (G) itaconate. * p <0.1 and ΔZ > |1| for the indicated comparison.

Figure S2. Plots for select metabolites altered at 6-9hpi in the indicated samples. (A) LysoPE (18:1), (B) LysoPE (18:2), (C) LysoPC (18:1), (D) LysoPC (18:2), (E) NAM, and (F) NMN. * p <0.1 and ΔZ > |1| for the indicated comparison.

Figure S3. Plots for select metabolites altered at 18-21hpi in the indicated samples. (A) Laurate, (B) 5-dodecenoate, (C) sarcosine, (D) proline, (E) β-alanine, (F) γ-glutamylthreonine (γ-Glu-Thr), and (G) γ-glutamylphenylalanine (γ-Glu-Phe). * p <0.1 and ΔZ > |1| for the indicated comparison.

Figure S4. Plots for polyamine metabolites in the indicated samples. (A) Putrescine, (B) N1-acetylspermidine (N1-Ac-Spd), and (C) N-acetylputrescine (N-Ac-Put). * p <0.1 and ΔZ > |1| for the indicated comparison.

Figure S5. Metabolite changes by stage in naïve larvae. (A) Volcano plot of metabolites that change with developmental stage. (B) Venn diagram comparing metabolite changes in *L. clavipes* infected larvae and with developmental stage.

**SUPPLEMENTAL TABLES**

Table S1. Statistical comparison of metabolites in *L. clavipes-*infected vs naïve larvae.

Table S2. Statistical comparison of metabolites in naïve larvae age-matched to 18-21hpi vs naïve larvae age-matched to 6-9hpi.

Table S3. Statistical comparison of metabolites in *msn>Pvr.λ* larvae vs genetic control larvae.
